# Chemotaxis and swarming in differentiated HL60 neutrophil-like cells

**DOI:** 10.1101/714345

**Authors:** Kehinde Adebayo Babatunde, Xiao Wang, Alex Hopke, Pierre-Yves Mantel, Daniel Irimia

## Abstract

Human leukemia cell line (HL-60) is an alternative to primary neutrophils in research studies. However, because HL-60 cells proliferate in an incompletely differentiated state, they must undergo differentiation before they acquire the functional properties of neutrophils. Here we provide evidence of swarming and chemotaxis in HL-60 neutrophil-like cells using precision microfluidic assays. We found that dimethyl sulfoxide (DMSO) differentiated HL-60 cells have larger size, increased length, and lower ability to squeeze through narrow channels compared to primary neutrophils. They migrate through tapered microfluidic channels slower than primary neutrophils, but faster than HL60s differentiated by other protocols, e.g. using all-trans retinoic acid. We found that differentiated HL-60 can swarm toward zymosan particle clusters, though they display disorganized migratory patterns and produce swarms of smaller size compared to primary neutrophils.

## INTRODUCTION

Neutrophils are immune cells that constitute an integral part of the first line of host defense against pathogenic infection and tissue injury ^1^. Upon pathogenic infection, neutrophils migrate rapidly from peripheral blood through the endothelial cell layer and into tissues, towards the site of infection ^2^. In response to infections and tissue injuries, neutrophils often display highly coordinated chemotaxis, accumulation, and clustering commonly known as “neutrophil swarming”, which is important in neutralizing infections and protecting healthy tissues. Better understanding of the neutrophil activities could eventually lead to interventions to enhance neutrophil efficacy against infections and protect tissues during inflammation. However, primary neutrophils are challenging to study directly because of short life span, donor variability, and low transcriptional activity that limits the use of common genetic approaches. Moreover, the isolation of neutrophils from whole blood is laborious and requires expensive isolation kits. Thus, relevant models are needed, and cell line-based models to substitute primary neutrophils would be highly useful.

The human leukemia cell line (HL-60) is a commonly used substitute cell line model to study neutrophil phenotypic functions. Several studies that have reported the use of HL-60 cells to study migration in neutrophils ^3 4 5 6^. These studies demonstrated migration in HL60 neutrophil like cells using assays like micropipette^6^, EZ-TAXIS assay ^6^, filter assay ^5^ and transwell assay ^7^. However, the information provided by those conventional assays are limited and seldom distinguishes the differences between migration and swarming of HL60 cells and primary neutrophils. Swarming activities *in vitro* and *in vivo* have so far only been observed using primary neutrophils ^8 9 10^. It is not known yet whether HL-60 cells could resemble the swarming behavior of primary neutrophils.

In this study, we employed microfluidic migration assays and a micropatterning array to compare the swarming and migratory ability of HL60s. We compared the migration of DMSO differentiated HL-60s (DdHL60), all-trans-retinoic acid (ATRA) HL-60s (AdHL60) and nutridoma supplemented DMSO differentiated HL-60s (nDdHL60) toward established chemo-attractants fMet-Leu-Phe (fMLP), Leukotriene B4 (LTB4), interleukin 8 (IL-8), and complement component 5a (C5a). We also compared the swarming ability of differentiated HL60s with that of primary neutrophils. We found that nutridoma differentiation enhances the migratory ability of HL60 neutrophil like cells compared to DMSO. Although both DdHL-60 and nDdHL-60 cells are capable of swarming, their behavior is quantitatively different than the swarming of primary neutrophils. Overall, our study suggests that nDdHL-60 cells are a good model for the chemotaxis studies in primary neutrophils, though differences in migration deformability, and persistence have to be taken into account.

## RESULTS

### Microfluidic assays for studying chemotaxis and swarming of dHL-60

The microfluidic device used to study chemotaxis consists of an array of a tapered channels, 500 m in length with cross-section area starting at (10 x 2 *μ*m^2^) and decreasing to (3 x 2 *μ*m^2^) connecting two opposite chambers namely; cell loading chambers and chemoattractant chambers^11^ (Fig. 1A) (Supplementary video. S1). A chemoattractant gradient is established along the tapered migration channels, from cell loading chambers to chemoattractant chambers (Supplementary video. S1). This device enables us to compare the chemotaxis and migration between HL-60 cell line model of neutrophil and primary neutrophil. To estimate number of chemotactic dHL-60 using tapered channel we calculated the percentage of cells migrating from the cell loading chamber through the tapered channel to the chemoattractant chamber (See method section for more details).

**Figure 1:**
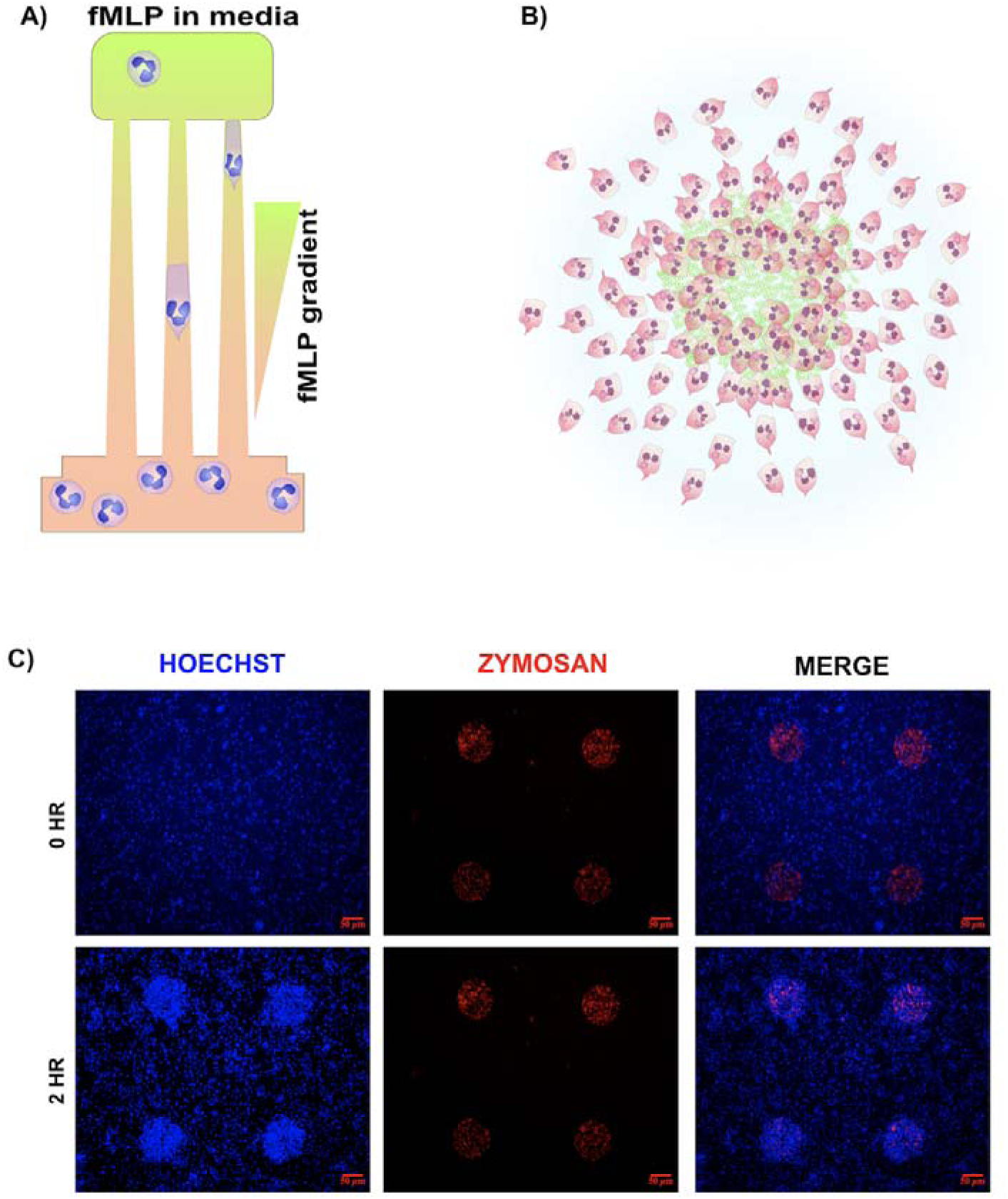
Schematic diagram of tapered channel and swarming assay. **A)** Tapered channel showing neutrophil cells migrating from the cell loading chamber (bottom) to the chambers loaded with chemoattractant fMLP (top). The tapered channel is approximately m in length and its cross-sectional area decreases from 20 μm to 6 μm B) Neutrophils migrate in an organized and directional manner toward zymosan particle cluster (green). C) Showing time lapse microscope image of dHL-60 swarming toward zymosan particle. First panel: Nucleus of dHL-60 stained with Hoechst at at 0 hr and 2hrs. Second panel: Zymosan particle clusters micro-patterned by poly-L-lysin and FITC-ZETAG solution. The size of the zymosan cluster spot is ∼140 μm^2^ and each spot is 1mm apart. Third panel: An array of dHL-60 swarm generated in the swarming generated using the swarming assay at 0 hr and 2hrs. Zymosan particle cluster target are covered by dHL-60 (blue).

To study swarm-like behaivour in dHL-60, we used a swarming assay designed by micro-pippetting a solution containing poly-L-lysin and FITC-ZETAG using a Polypico micro-dispensing machine. Next, we patterned zymosan particles clusters on the ZETAG spots on a plasma treated glass slide (Fig. 1A and 1C). Zymosan particle clusters were used as targets for both neutrophils and dHL-60 swarms (See method section for more details).

### DMSO stimulation enhances chemotaxis of dHL60 compared to AdHL60

We compared the migratory ability of HL60 neutrophils-like cells (dHL-60) after differentiation using different protocols to the migration ability of primary neutrophils. We exposed differentiated HL-60 cells to different chemoattractants and measured migration using a tapered microfluidic device with a cross-sectional area decreasing from 20 μm^2^ to 6 μm^2^ **(Fig. 1) (Supplementary video 1**). We found that 65%, 40.2%, and 2% of human neutrophils, DdHL-60, and AdHL-60 migrated in fMLP gradients, respectively (**Fig. 2**). The fraction of cells migrating was slightly larger in LTB4 and C5a gradients (70%, 25.7% and 8% for primary neutrophils, DdHL-60 and AdHL-60, respectively, for LTB4 and 75%, 27% and 4.3% for primary neutrophils, DdHL-60 and AdHL-60, respectively, for C5a). The fraction of cells migrating in IL8 gradients was significantly lower, at 17.6%, 16.3% and 5.5% for primary neutrophil, DdHL-60, and AdHL-60, respectively (**Fig. 2A**).

**Figure 2:**
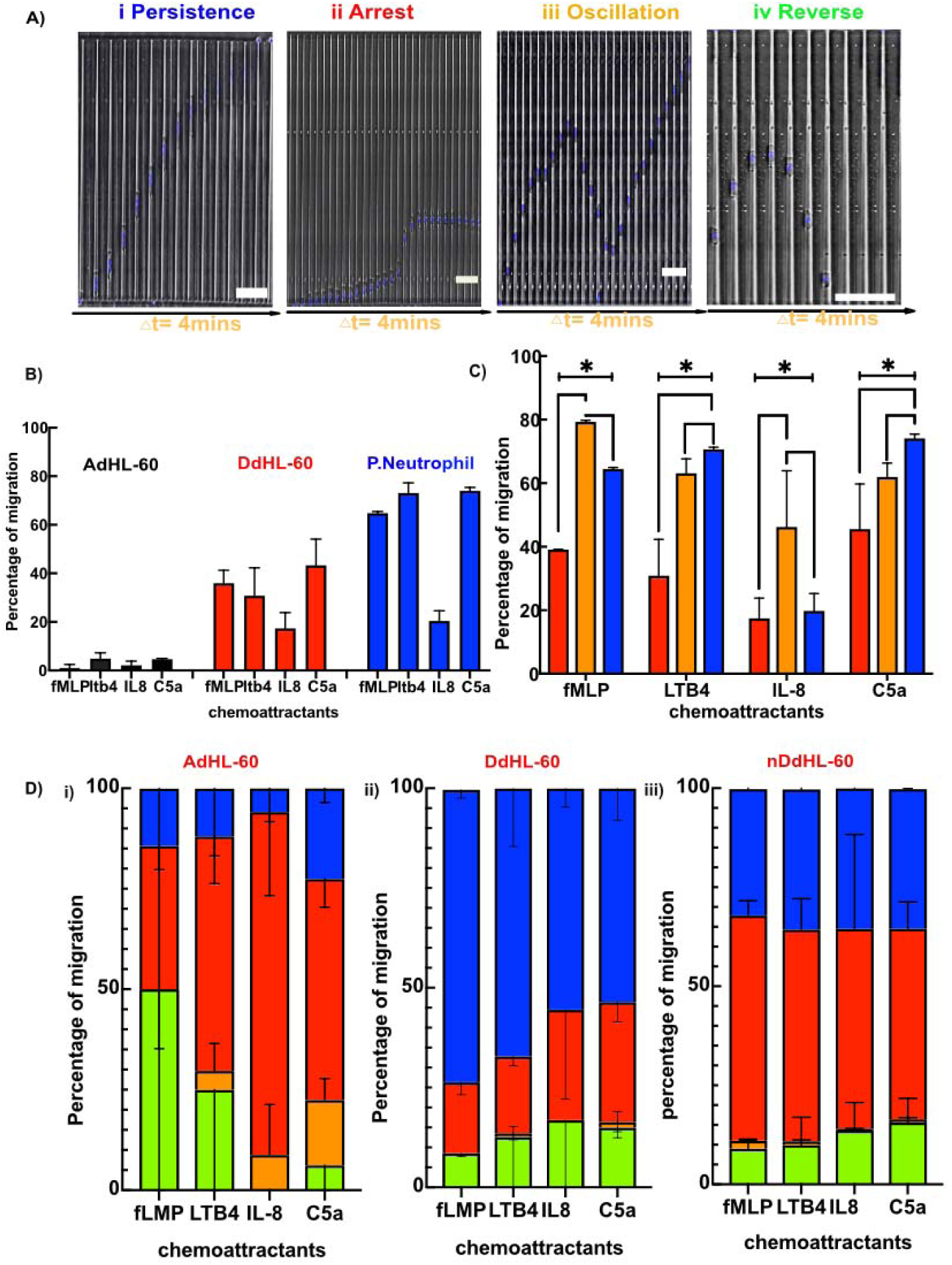
Migration of HL60 cells through tapered channels after differentiation using different protocols. A) Kymographs show the persistent (P), Arrest(A), Oscillation (O) and reverse (R) migration patterns. The nuclei of the HL60 cells are stained with Hoechst dye (blue). The time interval between two consecutive frames of the kymographs is 4 minutes. Scale bar is 25 µm. B) The percentage of migration in fMLP, LTB4, IL-8 and C5a gradients of ATRA (Black bars), DMSO (Red bars) and primary neutrophils (Blue bars). C) The percentage of migration of DdHL-60 (Red bars), nDdHL-60 (Orange bars), Primary neutrophils (Blue bars) towards each chemokine gradients (*P< 0.05, two-way ANOVA). D) Percentage of migratory patterns in i) DdHL-60 cells, ii) AdHL-60 cells and iii) nDdHL-60 cells. Persistence (Blue color), Arrest (Red color), Oscillation (Orange color) and Retrotaxis (Green color).

### Addition of nutridoma during DMSO-stimulated differentiataion of dHL60s enhances their chemotaxis

Nutridoma, a serum-free supplement, has been reported to improve the differentiation of PLB-985 cells while not compromising the viability of the cells ^12 13^. To test how serum concentration and nutridoma affect differentiation in HL-60 cells, we checked the chemotactic ability of HL-60 cells upon differentiation in media containing 1.3% DMSO, 2% fetal bovine serum (FBS) and 2% nutridoma. Replacing serum with nutridoma increased the percentage of migrated cells to 78, 41, 34, and 59% in fMLP, LTB4, IL-8, and C5a gradient compared to 40, 36, 20, and 45% for DMSO alone, respectively (**Fig. 2B**). Together, our data showed that nDdHL60 cells and primary neutrophils are comparable in terms of migratory ability.

### Characterizations of migratory patterns of dHL-60 using tapered channel microfluidic device

Using the tapered channel, we identified four migration behaviors in dHL-60 cells, similar to those observed in primary neutrophils ^11^ including persistent migration (P), arrest (A), oscillation (O), and retro-taxis (R) (**Fig. 2C**) (**Supplementary video 1**). Persistent migration indicates neutrophils that migrated through the tapered channels without changing directions from the cell loading chamber to the chemoattractant chamber. Arrest describes neutrophils that migrated into the tapered channels but got trapped in the channels. Oscillation indicates neutrophils that change migration direction more than two times within the tapered channel. Retro-taxis describes neutrophils that migrated into the tapered channel but migrated back into the cell-loading chamber. To calculate the percentage of each migratory patterns, we counted the number of cells exhibiting a specific pattern compared to the total number of migrated cells (See method section for more details). These migratory patterns are chemokine-dependent. 73, 66, 63 and 55% of DdHL-60 neutrophils and 25, 12, 30 and 11% of the AdHl-60 neutrophils migrated persistently in fMLP, LTB4, C5a and IL-8 gradients respectively (**Fig. 2Di, Fig. 2Dii**) compared to 84, 76 and 70% of primary neutrophils migrated persistently in fMLP, LTB4 and IL-8 respectively^11^. In fMLP gradient, 18.5, 7.5, and 0% of the DdHL-60 and 58.5, 24.4 and 4.9% of AdHL-60 neutrophils showed arrest, retro-taxis, and oscillations, respectively (**Fig. 2Di, Fig. 2Dii**). In contrast, 7, 7, and 2% of primary neutrophils showed arrest, retrotaxis and oscillation in fMLP gradient respectively ^11^. The migration patterns of AdHL-60 included larger percentage of arrest and retro-taxis compared to DdHL-60 (**Fig. 2Dii**). Surprisingly, the presence of nutridoma in the differentiation media markedly changed the migratory patterns of nDdHL60 cells, which display frequent arrest and retrotaxis migratory patterns.. We observed that 36.4%, 35.4%, 23.6% and 35.2% migrated persistently in fMLP, LTB4, IL-8 and C5a gradient respectively compared to DMSO alone (**Fig. 2Diii**). In addition, about 65.5%, 64.7%, 76.4% and 64.8% of nDdHL-60 demonstrated more arrest migratory pattern in all chemokine gradient compared to DMSO alone (**Fig. 2Diii**).

The tapered microchannels enable specific measurements of the cross-section (CS) at which A, O, and R patterns occur althogether in AdHL60, DdHL60 and nDdHL60 (**Fig. 3A-C**). Our results demonstrate that the average CSs of AdHL60 are 10, 18, 17 and 15 µm^2^ in fMLP and LTB4, C5a and IL-8 gradients respectively (**Fig. 3A**), DdHL-60 are at 10, 10.5, 11 and 10.5 µm^2^ in fMLP and LTB4, C5a and IL-8 gradients respectively (**Fig. 3B**) and the average CSs of nDdHL-60 are at 13.5, 13.5, 13 and 13.5 µm^2^ in fMLP and LTB4, C5a and IL-8 gradients respectively (**Fig. 3C**). While the average CSs of primary neutrophils are at 10.5, 9.2 and 10 µm^2^ in fMLP, LTB4 and IL-8 gradient respectively ^11^.

**Figure 3:**
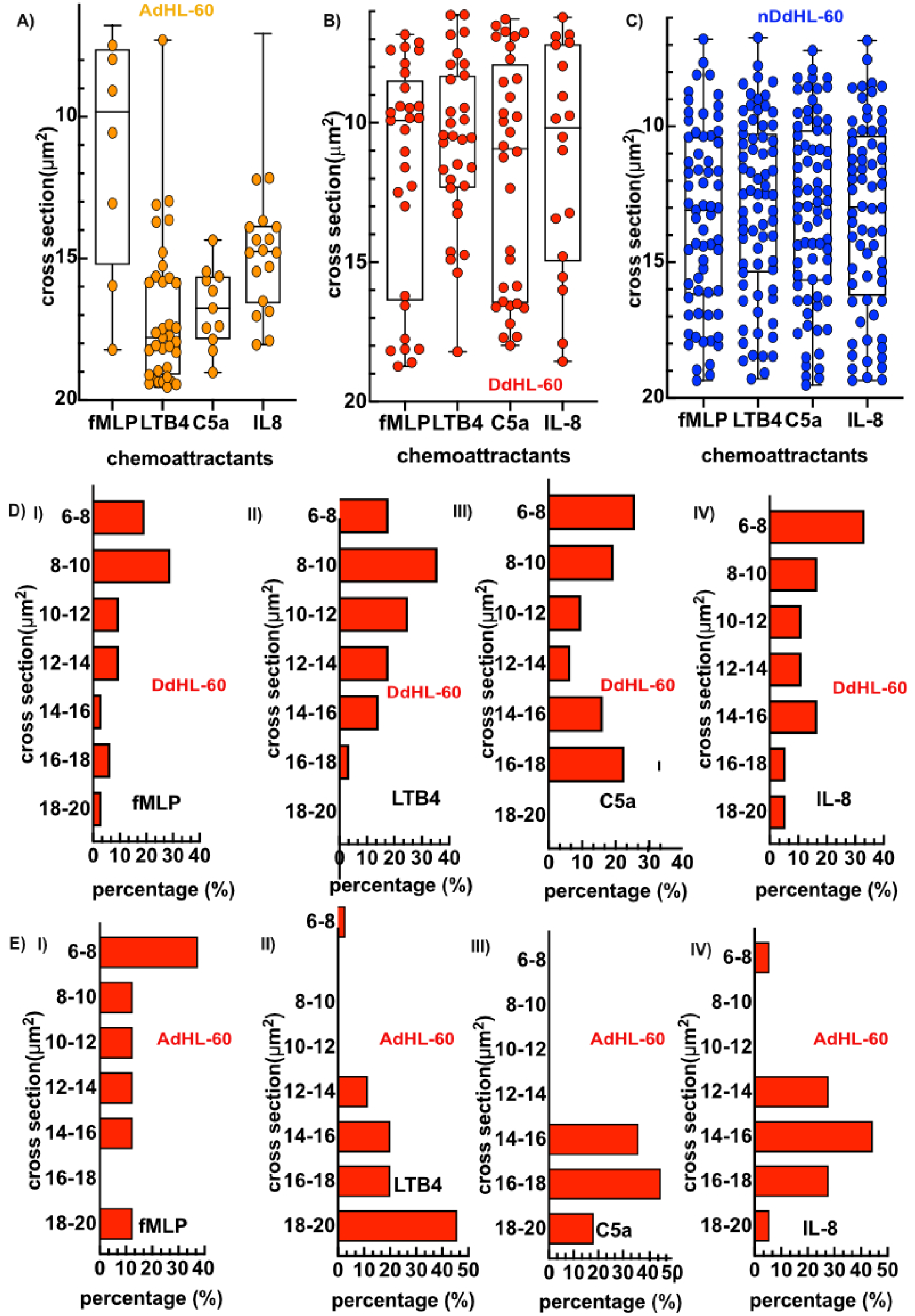
The differentiation protocol alters the ability of HL60 cells to traverse narrow strictures. A) Critical cross section where Arrest, Oscillation and Retrotaxis occur in A) AdHL-60(Orange) B) DdHL-60(Red) and C) nDdHL-60(Blue) in tapered straight channel in chemokine gradients of fMLP, LTB4, IL-8 and C5a. Histograms of Arrest, Oscillation and Retrotaxis critical sections in D) DdHL-60 cells and E) AdHL-60 in different chemokine gradients.

The histograms of the critical cross-section show that larger percentage of A, O, R patterns in DdHL-60 cells occur only at a cross-section of lower than 10 µm^2^ (**Fig. 3Di-iv**) similar to what was observed in primary neutrophils ^11^. The histograms of the critical cross-section show that A, O, R patterns in AdHL-60 cells are less dispersed (**Fig. 3Ei-iv**). In summary, our data demonstrated that mechanical constriction in the tapered channel during chemotaxing interfered and altered the migratory response of dHL-60 to chemokine gradients.

### Morphological comparison of chemotactic dHL-60 and primary neutrophils in confined microfluidic channels

Our data showed that dHL60 are larger and longer than primary neutrophils during chemotaxis. We observed that migrating HL-60 neutrophil-like cells(dHL-60) display a broad leading edge toward the chemokine and a rounded tail end (**Fig. 4A**). When the cells advance through the tapered channels towards the smaller cross sections, their length gradually increases. The length of DdHL-60 cells, nDdHL-60 cells and primary neutrophils increases from ∼38 to ∼78 µm, ∼30 to ∼65 µm and from ∼25 to ∼65 µm respectively when migrating through the tapered micro-channel toward the chemokine (**Fig. 4B**). We observed that dHL60 cells migrated slower in all chemokine gradients compared to primary neutrophils. We found that DdHL-60 cells reach 24 (6.6 ± 6.6) and∼15 (5.8 ± 6.3) µm/min velocity in fMLP and LTB4 gradients, respectively (n = 10) (**Fig. 4C and 4D**). Surprisingly, nDdHL-60 cells are slower compared to DMSO alone as they migrate at a velocity of 15 (4.8 ± 7.3) and 12 (4.5± 11.5) µm/min in fMLP and LTB4 gradients, respectively (n=10) (**Fig. 4C and D**). The velocities of both DdHL-60 and nDdHL-60 cells in fMLP gradient decreased with decreasing cross section (**Fig. 4C**). The highest velocity of DdHL60 cells was recorded at 13-15 µm^2^ cross-section in LTB4 gradients. While the highest velocity of nDdHL-60 was recorded at 18-20 µm^2^ cross-section in LTB4 gradients (**Fig. 4D**). DdHL-60 and nDdHL-60 cells are both slower than primary neutrophils, which can reach velocities as high as 35 (13.6 ± 16.8) and 42 (17 ± 16.8) µm/min in fMLP and LTB4, respectively (n=10) (**Fig. 4C and 4D**). The highest velocity in primary neutrophils was recorded at a cross-sectional area between 16-18 and 14-16 µm^2^ in fMLP and LTB4 gradients, respectively (**Fig. 4C and 4D**).

**Figure 4:**
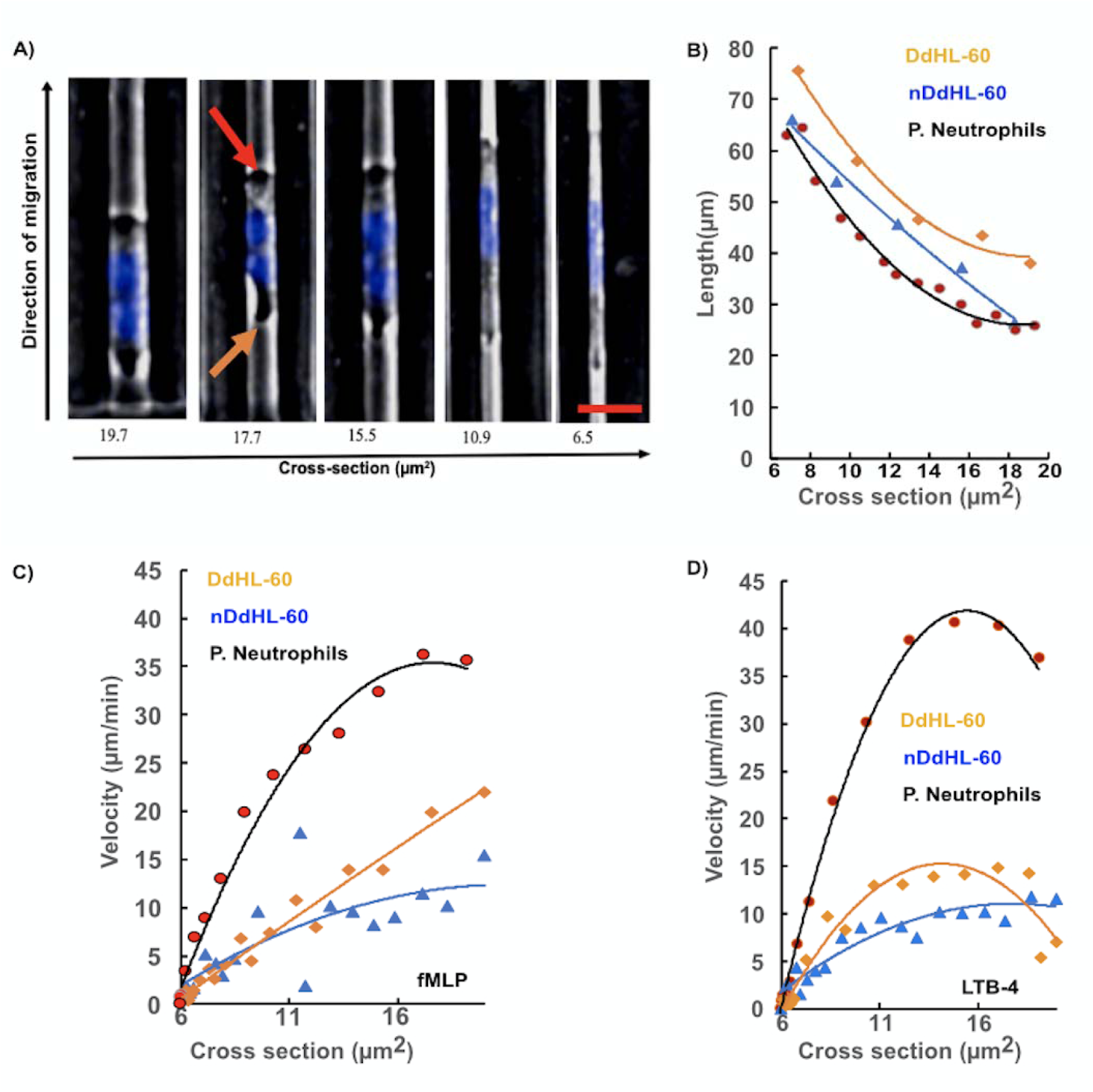
Comparison of cell length, velocity, and velocity changes between dHL-60 cells and primary neutrophils across the different cross-sections of the tapered micro-channel. A) The morphology of dHL-60 cells across changes during the migration through progressively narrow channels. In the larger section of the channels, HL60 cells display a broad leading edge (Red arrow) and a rounded tail end (uropod) (Orange arrow). The nucleus (blue) of dHL-60 neutrophil deforms to accommodate the migration through the narrowest section of the channels. The leading edge and uropod are no longer distinguishable. HL60 nucleus is stained with Hoechst dye. The scale bar is 30 µm. B) Change in length across different cross-section of the tapered micro-channel DdHL-60 cells (Orange line), nDdHL-60 cells (Blue line) and primary neutrophil (Black line) C) Change in velocity in fMLP across different cross-section of the tapered micro-channel DdHL-60 cells (Orange line), nDdHL-60 cells (Blue line) and primary neutrophil (Black line). D) Change in velocity in LTB4 across different cross-section of the tapered micro-channel DdHL-60 cells (Orange line), nDdHL-60 cells (Blue line) and primary neutrophil (Black line)

### dHL-60 is less directional and organized than primary neutrophil in swarm-like behavior

Next, we compared the swarming ability of human neutrophils with that of the differentiated cell lines, DdHL-60, and nDdHL-60. We observed that human neutrophils swarm efficiently toward zymosan particle clusters with three distinct phases. Swarming starts with random migration on the surface (*scouting phase*-5 mins **Fig. 5A**). After the first neutrophil interacts with the cluster, the number of migrating neutrophils towards the zymosan cluster increases rapidly (*growing phase- 10-15 mins* **Fig. 5A**). Swarms reach their peak size 60-120 minutes later, after which the size remains stable (*stabilization phase*-**Fig. 5A**) (**Supplementary video 2)**. We observed qualitatively similar aggregation dynamics in both DdHL-60 and nDdHL-60 cells. Aggregation starts with the cells moving randomly on the surface of the zymosan particle clusters (*scouting phase*-5mins **Fig. 5A**). Within minutes of interaction with the zymosan particle, an increasing number of dHL-60 cells migrate in the direction of the zymosan particle clusters (*growing phase*- 10-30 mins **Fig. 5A**). A fast growth is followed by slower growth. Swarms in nDdHL-60 cells reach their peak size after 60-120 minutes, after which the size of the aggregates decreases because cells leave the aggregate (**Fig. 5A**). During the stabilization phase, dHL-60 cells migrate in and out of the swarms (**Fig. 5A**) (**Supplementary video 3 and 4**).

**Figure 5:**
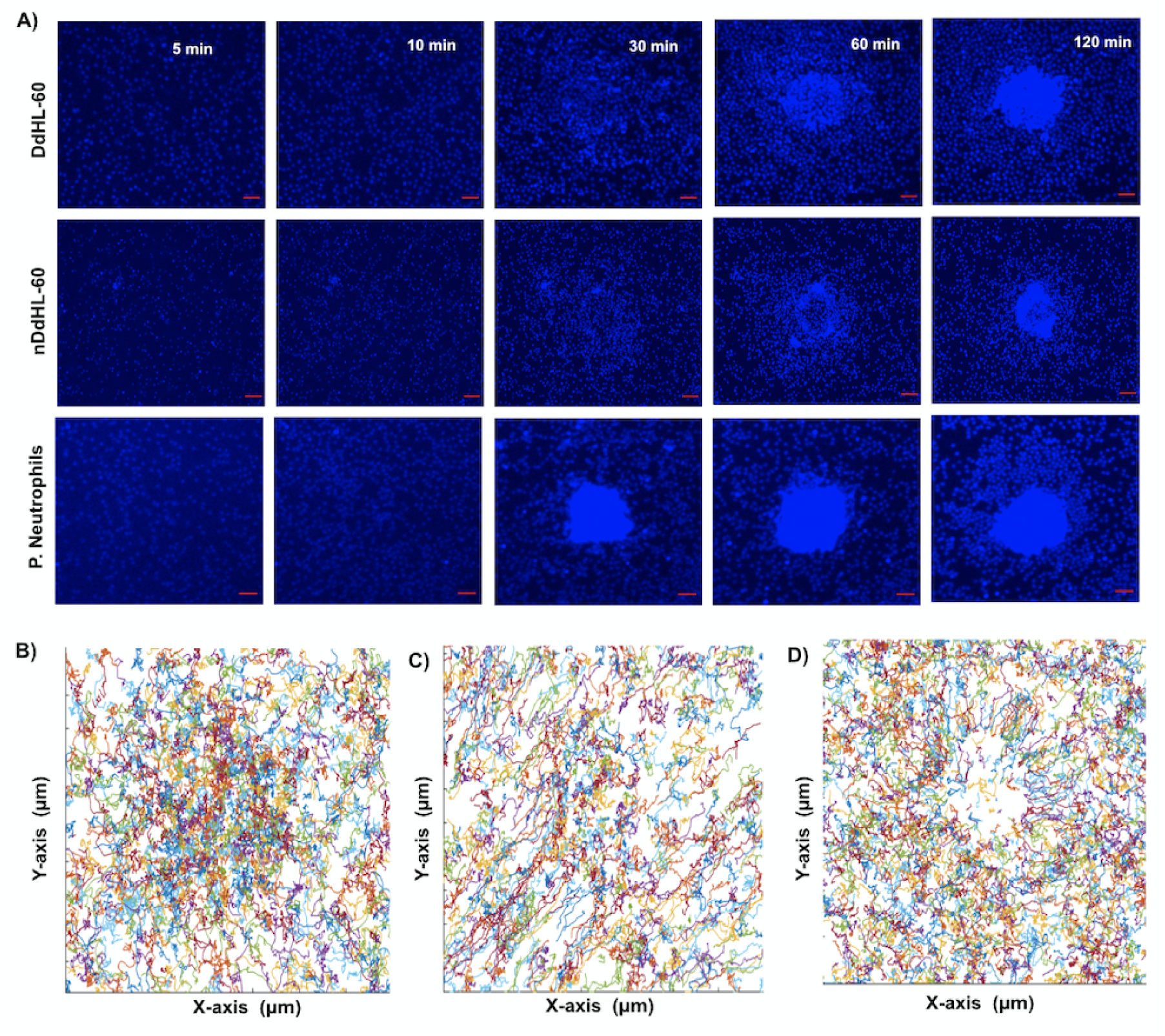
Swarming and aggregation of DdHL-60, nDdHL-60, and primary neutrophils on zymosan-cluster targets. A) Sequential images showing swarm formation in DdHL-60, nDdHL-60 and primary neutrophils (blue) around patterned zymosan particle clusters at 5mins, 10mins, 30mins, 60mins and 120 mins (scale bar: 25 µm). Moving B) DdHL-60 C) nDdHL-60 D) primary neutrophils display random and precise track trajectories respectively toward centered zymosan particle. Figure shows representative experiments of N= 4.

A quantitative analysis of neutrophil swarming showed significant differences between dHL-60 and human neutrophil swarming. We found that the radial trajectories displayed by dHL-60 cells are more often random and less organized (**Fig. 5B and 5C**) compared to swarming primary neutrophils (**Fig. 5D**, **Fig. 6Ai, Supplementary video 2)**. Both DdHL-60 and nDdHL-60 cells show increasingly directional migration toward the zymosan clusters at short distances around the zymosan clusters (**Fig. 6Aii and iii**). The migratory speed increases steadily between 10-100 minutes towards the zymosan particle cluster in primary neutrophils (**Fig. 6Bi**). However, nDdHL-60 cells show only slight increase in migratory speed toward zymosan particle cluster (**Fig. 6Biii**). The swarm area around zymosan clusters during the stabilization phase was larger for primary neutrophils compared to dHL60 cells, ∼40,000 μm^2^ vs. 25,000 μm^2^, respectively (**Fig. 6C**). The mean swarm area around zymosan clusters was ∼25% larger for primary neutrophils compared to dHL-60 cell, at 25,000 μm^2^ vs. 20,000 μm^2^, respectively (**Fig. 6D**).

**Figure 6:**
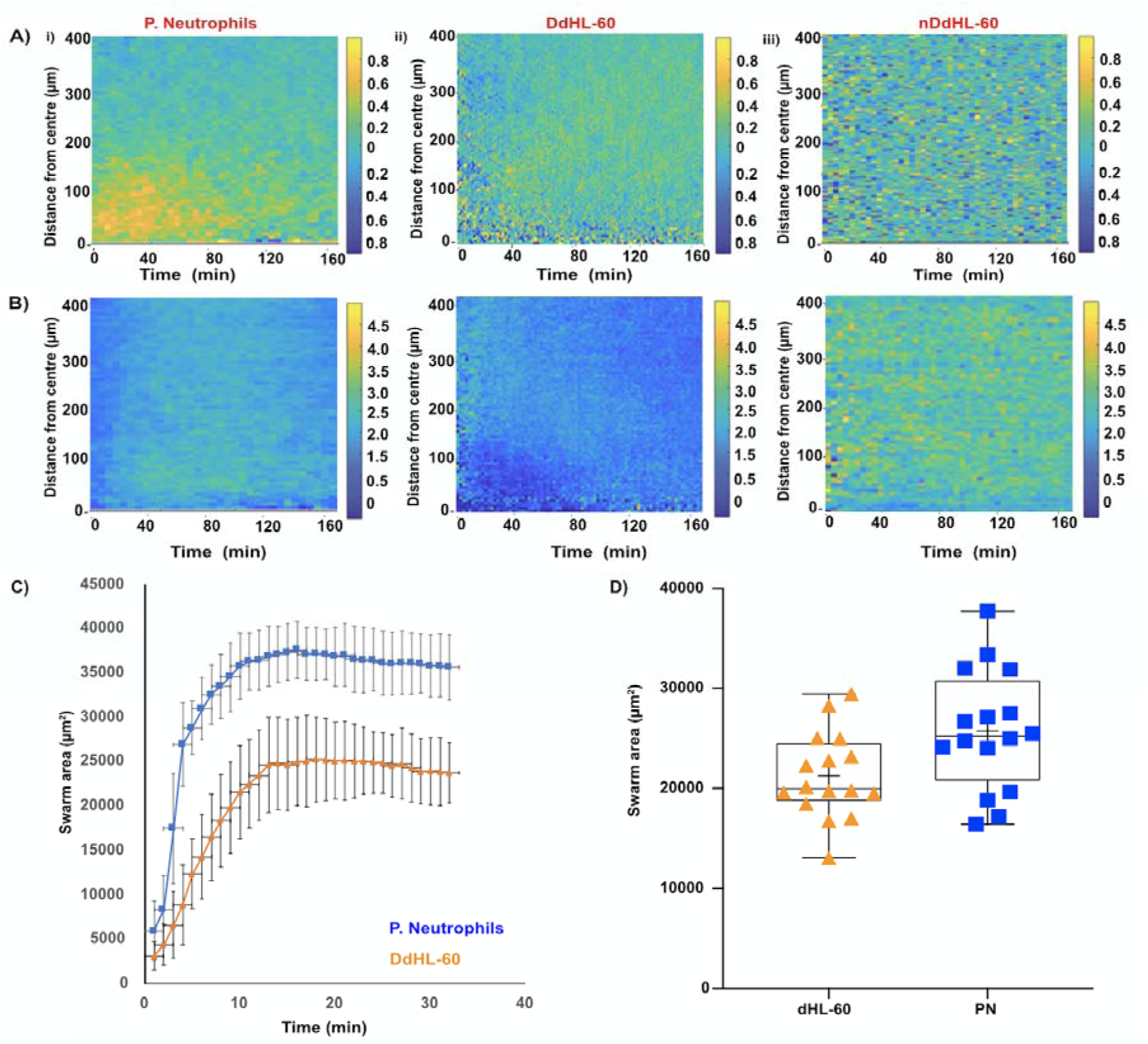
Quantification of primary neutrophil and HL60 swarming. A) Chemotactic index (CI) and speed over time, towards zymosan particle clusters in i) primary neutrophils ii) DdHL-60 iii) nDdHL-60. B) The speed of migration increases with time towards zymosan particle clusters. (Scale bar: CI above 0.8 indicates the cells that are chemotaxing toward zymosan particle. Speed (µm/min) = 4.5 means cell moving toward zymosan particle). C) Primary neutrophils (blue curve) accumulation on targets is fast and form a bigger swarm and dHL-60 neutrophil (orange curve) aggregation proceeds fast and then continues slower over time. D) Comparison of area of swarm formed between primary neutrophils (PN) and dHL60 (HL-60). Error bars represent standard deviations for these measurements. Figure 5 shows representative image of experiment of N= 30 swarm spots.

### Blocking LTB-4 receptors alters dHL-60 swarm-like behavior

To verify that the aggregation of HL60 cells is phenotypically similar to swarming, we tested the role of LTB4 release in the positive feedback loop driving the rapid accumulation of neutrophils during the growth phases of the swarms. We compared the aggregating of HL60 cells in the presence of BLT1 and BLT2 receptors antagonists and LTB4 synthesis inhibitor. We found that in the presence of BLT1 and BLT2 receptor antagonists LY255283, U75302 there was significant delay in the initiation and a reduction in the swarm size (**Fig. 7A and 7B**) (**Supplementary video 5 and 6**). The inhibition of the LTB4 pathway by the MK886 inhibitor alters mainly the final size, with little impact on the initiation, suggesting the release of pre-stored LTB4 during the early stage of swarming (**Fig. 7A and 7B**) (**Supplementary video 5 and 6**). The swarm area around zymosan clusters during the stabilization phase was ∼2 times larger in control compared to drug treated cells (∼25000 μm^2^ vs. 17000μm^2^ vs. 15000 μm^2^, for control vs. BLT1&2 vs. MK-886 treated cells) (**Fig. 7B**).

**Figure 7:**
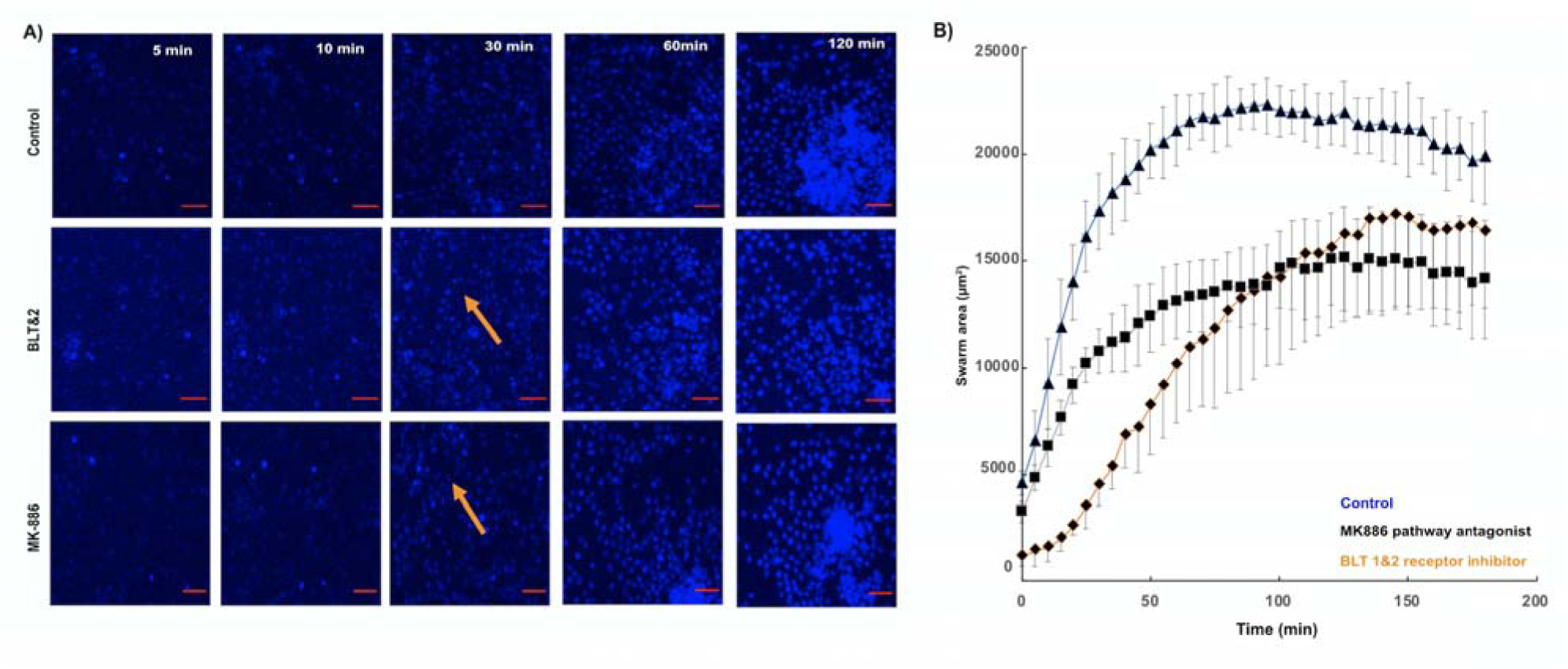
Disrupted migration of DdHL-60 towards swarm in the the presence of BLT1 receptor antagonist and LTB4 synthesis inhibitor. A) Sequential images showing swarm formation in Control, BLT1&2 receptor antagonist and LTB4 pathway antagonist around patterned zymosan particle clusters at 5mins, 10mins, 30mins, 60mins and 120 mins (scale bar 25 µm). Orange arrows showing delay in initiation. B) dHL-60 form smaller swarms that grow slower in the presence of BLT1 antagonists and LTB4 pathway inhibitors. (Blue (triangle) curve: Control, Black (square) curve:LTB4 pathway antagonist MK-886, Orange(diamond) curve: BLT1&2 receptor antagonist). Each group was measured in four separate spots in a field of view. Error bars represent standard deviations for these measurements. Figure shows representative experiments of N= 36 fields of view (4 spots per field).

## DISCUSSION

Neutrophil swarming has been described as a crucial process of neutrophil tissue response needed to specifically regulate tissue protection and destruction during several inflammatory diseases ^14^. Here, we report that HL60 are capable of swarming, qualitatively similar to primary neutrophils. Previously, neutrophil swarming has been shown in zebrafish larvae ^15 16^ and mouse tissues ^10 17 9^. Human neutrophil swarming has also been observed *in vitro* ^18 8^. Although dHL-60 cells can synthesize LTB4 and also possess LTB4 receptors^19 20 13^, the swarming of HL-60 has not been demonstrated. By micropatterning zymosan particles, we reported the first evidence of swarm-like behavior in dHL-60 cells.

Swarming in primary neutrophils is thought to occur in three phases: (1) early neutrophil migration(*scouting*) followed by (2) amplification of migratory response (*amplification*) and finally (3) clustering at the site of infection(*stabilization*) ^21^. The striking feature of dHL-60 cells behavior during swarming is the display of all three phases of swarming (scouting, amplification, and stabilization phases) ^8 10^. However, despite the qualitative similarities, significant differences exist. The swarm size for similar targets is smaller and the trajectories of cells joining the swarms appear more disorganized for differentiated HL60 cells compared to primary neutrophils.

The differences in chemotaxis (migration fraction, migratory velocity, and directionality) between dHL-60 and primary neutrophils may contribute to the differences observed in the swarming assay. A lower percentage of dHL60 cells migrate towards chemoattractants compared to primary neutrophils and which may be responsible for the observed smaller swarm size in dHL60. Moreover, our study demonstrates that dHL60 cells migrate slower than primary neutrophils^10 17 9^. More importantly, the third phase of swarming is characterized by recruitment of distant neutrophils to an infection site, driven by expression of high-affinity receptor for LTB4 (LTB4R1) on neutrophils to maintain persistent and long-term neutrophil recruitment ^10^. Our data demonstrates that the observed swarm-like behavior in dHL-60 cells is LTB4 dependent, similar to the findings in primary neutrophils (^8^. Our chemotaxis data shows that a lower percentage of dHL-60 cells migrated toward LTB4 with more arrest and retrotaxis which may be the cause of the smaller swarm size observed in dHL60 cells.

In summary, our study provides first evidence for swarming behavior in dHL-60 cells. The dHL-60 swarms are smaller than the primary neutrophils. While the migration speed is comparable, the observed differences may be due to the differences in the expression of LTB4 on the differentiated dHL-60. Deficits in the mechanisms of intracellular communication or a combination of these factors may also be involved. Our study, therefore, suggests that DdHL-60 and nDdHL-60 could be useful models for the study of neutrophil chemotaxis and swarming with further optimizations.

## MATERIALS AND METHODS

### HL-60 culture under DMSO and ATRA exposure

HL-60 cells (CCL-240, ATCC, Manassas, VA) were either cultured in RPMI 1640 containing with 10% heat-inactivated fetal bovine serum (FBS), 5mL antibiotics or RPMI 1640 containing 2% FBS, 2% Nutridoma and 5mL Antibiotics at 37°C in 5% CO_2_. Cells were passaged when a maximal density of 1–2 million cells per mL was reached usually every 2-3 days. Passage were done in a total volume of 10 mL RPMI 1640 + 10%FBS media, in 75 cm^2^ cell culture flasks. HL60 cells were differentiated with 1.3% v/v DMSO for 6 days with antibiotics or 5 μM of ATRA for 5 days without media change. For nuclei staining HL-60 neutrophil-like cells were treated with 0.5 μg/ml Hoechst 33342 (Life Technologies) for 5 minutes at 37 °C and washed in PBS once. During migration assays, neutrophils like cells were suspended in appropriate RPMI media

### Neutrophil isolation

Human blood samples from healthy donors (aged 18 years and older) were purchased from Research Blood Components, LLC. Human neutrophils were isolated within 1h after drawn using the human neutrophil direct isolation kit (STEMcell Technologies, Vancouver, Canada) following the manufacturer’s protocol. After isolation, neutrophils were stained with Hoechst 33342 trihydrochloride dye (Life Technologies). Stained neutrophils were then suspended in RPMI 1640 media containing 20% FBS (Thermo Fisher Scientific) at a concentration of 2 × 10^7^ cells/ml in the chemotaxis device or 2.5 x 10^6^ cells/mL in the swarming assay.

### Neutrophil inhibition

LY255283, U75302, and MK-886 were dissolved in DMSO. Each compound was re-suspended in cell culture media at a concentration of 20 µM for LY255283 and U75302 and 400nM for MK-886 pathway inhibitor. For BLT1&2 receptors inhibition, dHL-60 were incubated in LY255283, U75302 for 30 minutes and for LTB4 synthesis inhibition, dHL-60 were incubated in MK-886 for 30 minutes before adding on the swarming assay.

### Device fabrication

The microfluidic devices were fabricated as described by Wang and colleagues using standard soft lithography ^11^. Briefly, two-layer master mold in negative photoresist (SU-8, Microchem, Newton, MA) were fabricated on a 4-inch silicon wafer. The first layer was 2 *μ*m thin containing the patterns of the tapered migration channels. The second layer was 75 *μ*m thick and consists of cell-loading channels (CLC) and chemokine chambers. A ratio of 10:1 PDMS base and curing agent were mixed, cast on the master mold, and degassed thoroughly (PDMS, Sylgard, 184, Elsworth Adhesives, Wilmington, MA). We transferred the wafer into an oven at 65◦C to cure overnight. After curing, we peeled off the PDMS layer from the wafer and cut out individual devices using a scalpel. We punched the inlets and outlets of the devices using a 0.75 mm diameter biopsy puncher (Harris Uni-Core, Ted Pella) and irreversibly bonded them to a glass-bottom multi-well plate (MatTek Co., Ashland, MA).

To prepare the glass slides for swarming assay, plasma treated glass slides (Fisher brand Double Frosted Microscope Slides, Fisher Scientific, Waltham, MA, USA) were micro-patterned with a solution containing poly-L-lysin and FITC-ZETAG (1.6mg/ml) using a Polypico micro-dispensing machine. Zymosan particle clusters were used as targets for neutrophils swarms. A solution of 0.5 mg/mL zymosan particles in ultra-pure water (Gibco, life technologies, USA) was prepared and sonicated for 10 minutes before pipetting onto the glass slide and and was allowed to adhere for 10mins on a hot plate. Excess zymosan particles was then washed thrice with PBS and stored at room temperature. Prior to experiment, glass slides were placed in an open well chamber. For HL-60 neutrophil-like cells, the chambers were coated with 50ug/ml of fibronectin for 1 hour at 37°C to improve migration of the cells.

### Microfluidic devices to study neutrophil chemotaxis

The microfluidic device used for this study was designed as described by ^11^ and it consists of an array of tapered channels, with a cross-sectional area of 20 *μ*m^2^ at the cell loading chamber end to 6 *μ*m^2^ at the chemo-attractant chamber end. The tapered channels are 500 *μ*m in length and connect the cell-loading chamber (CLC) to several chemoattractant chambers. A chemoattractant gradient that increases toward the chemoattractant chamber is established along the tapered migration channels. This device enables us to compare the chemotaxis and migration between HL-60 cell line model of neutrophil and primary neutrophil. Using time-lapse imaging and automatic cell tracking on image J software, we were able to effectively compare the active migration patterns between a single DdHL-60 neutrophil-like cell and a single primary neutrophil with high spatial and temporal resolutions.

### Device set-up and cell loading

fMLP (Sigma-Aldrich), LTB4 (Cayman Chemical, Ann Arbor, Michigan, USA), C5a and IL-8 (Cayman Chemical, Ann Arbor, Michigan, USA) were diluted in RPMI to a working concentration of 100 nM. C5a was diluted in RPMI without FBS to working concentrations of 100 nM. Five microliters of the chemokine solution were pipetted into each device. The well plate was then placed in a desiccator under vacuum for 15 min. The well plate was then taken out from the desiccator for 15 min until the devices were filled with the solution. Three milliliters of media (RPMI + 10% FBS) was then added to each well to cover the devices. Each device was then washed by pipetting 10 *μ*l media through the inlet. Finally, about 5 x10^4^ HL-60 neutrophil-like cells were pipetted into each device. Time-lapse images of neutrophil migration were captured at 10°ø magnification using a fully automated Nikon TiE microscope (Micro Device Instruments). The microscope is equipped with a bio chamber heated at 37◦C and 5% CO2.

### Analysis of differentiated HL-60 neutrophil-like cells (dHL-60) migration

We used Track-mate module in Fiji ImageJ (ImageJ, NIH) to track and analyze cell trajectories automatically. We identified four migration behaviors, including persistent migration (P), arrest (A), oscillation (O), and retro-taxis (R). Persistent migration indicates neutrophils that migrated through the channels without changing directions. Arrest describes neutrophils that are trapped in the channels. Oscillation indicates neutrophils that change migration direction more than two times. Retro-taxis describes neutrophils that migrated back to the cell-loading channel. Percentage of migration was calculated as thus: (N_mc_)**/**(N_CLC_)*100. Where N_mc_ is total number of migrated cells in the tapered channel and N_CLC_ is total number of cells in the cell loading chamber. Percentage of each migration pattern was calculated as thus: (N_mp_)**/**(N_mc_)*100. Where N_mp_ is number of cells demonstrating a specific migration pattern and N_mc_ is total number of migrated cells in the tapered channel.

### Swarm size measurement

Changes in swarm size over time were estimated using track mate plugin in Image J. The cell-occupied area was measured from the Hoechst labeled HL-60 neutrophil-like cells and primary neutrophil using filter and suitable threshold (otsu and Huang) on image J.

### Chemotactic and tracking analysis

To quantify the directional radial migration during swarming, we measured the distance of neutrophils to the zymosan spot and plotted the changes in this distance for individual tracks over time. The instantaneous chemotactic index (CI) at time *t* was then calculated as: CI(t)=−R′(t)/X′(t) where: R′(t)=∂/∂t x−z is the rate of change of the distance between the cell’s position *x* and the zymosan particle cluster position *z*. Before chemotaxis, analysis the cell track positions *x* was determined using track mate plugin in Image J. Clusters of zymosan particles were segmented by Huang thresholding followed by selection of the largest connected component, and *z* was set to be the centroid of this object. Cellular migration speed (spline smoothed) is calculated as: X′(t)= ∂x/∂t.

### Statistical analysis

Statistical significance of the differences between multiple data groups were tested using two-way Analysis of Variance (ANOVA) in GraphPad Prism (GraphPad Software). Within ANOVA, significance between two sets of data was further analyzed using two-tailed **t**-tests. All the box plots consist of a median line, the mean value (the cross), and whiskers from minimum value to maximum value.

## ACKNOWLEDGMENTS

We thank Miss. Jiseon Min and Dr. Amir Ariel of John A. Paulson, School of Engineering and Applied Sciences, Harvard University for their help with the cell traffic analysis during swarming. This work was supported by funds from the National Institutes of Health (GM092804 and EB002503). Kehinde Babatunde was supported by a Swiss National Funding for Doctoral Mobility fellowship (P1FRP3_181378).

## AUTHOR CONTRIBUTIONS

P.M and D. I. designed research. X.W. and K.A.B. designed and prepared microfluidic devices. K.A.B., A.H., and X.W conducted experiments. K.A.B., X.W. and D.I. analyzed results. K.A.B., X.W., P.M. and D.I. prepared the manuscript, with significant contributions from all authors.

