## Supplementary material for "Chemotaxis and swarming in differentiated HL60 neutrophil-like cells": Captions

**SUPPLEMENTARY VIDEOS**.

**Supplementary Video 1.** **dHL-60 neutrophil like cells moving through tapered channels**. dHL-60 neutrophil like cells migrate from the cell loading chamber (bottom) to the chambers loaded with chemoattractant fMLP (top). The tapered channel is approximately 500 μm in length and its cross-sectional area decreases from 20 μm^2^ to 6 μm^2^. Four distinct migratory patterns of dHL-60 neutrophil like cells are outlined. The green circle shows a persistent migratory pattern, red shows arrested migratory pattern, yellow indicates retrotaxis, and orange indicates oscillation migratory pattern. The time interval between frames is 4 minutes. Scale bar is 50 μm. The nucleus of the dHL-60 cells is stained with Hoechst dye (blue).

**Supplementary Video 2. Primary neutrophil swarming on clusters of zymosan particles**. Primary neutrophils (blue - nucleus stained with Hoechst dye) swarm around four zymosan particle-cluster spots (140 μm in diameter and 500 μm apart). The spots are identified by broken red circles. The three distinct phases of swarming are outlined: a scouting phase in the first 5-10 mins followed by a growing/amplification phase and a stabilization phase at approx. 60 mins after loading. Isolated neutrophils are loaded on the swarming assay device at a concentration of 2.5 x 10^6^ cells/mL. The time interval between frames is 5 minutes. Scale bar is 50 μm.

**Supplementary Video 3. dHL-60 swarming on clusters of zymosan particles**. dHL-60 cells (blue - nucleus stained with Hoechst dye) swarm around four zymosan particle-cluster spots (140 μm in diameter and 500 μm apart). The spots are identified by broken red circles. The three distinct phases of swarming are outlined: a scouting phase in the first 5-10 mins followed by a growing/amplification phase and a stabilization phase at approx. 60 mins after loading. The migration of dHL60 during swarming is less organized and less directional compared to primary neutrophils. Isolated neutrophils are loaded on the swarming assay device at a concentration of 2.5 x 10^6^ cells/mL. The time interval between frames is 5 minutes. Scale bar is 50 μm.

**Supplementary Video 4: dHL-60 swarming on clusters of zymosan particles**. dHL-60 cells (blue - nucleus stained with Hoechst dye) swarm around four zymosan particle-cluster spots (140 μm in diameter and 1mm apart). The spots are identified by broken red circles. The three distinct phases of swarming are outlined: a scouting phase in the first 5-10 mins followed by a growing/amplification phase and a stabilization phase at approx. 60 mins after loading. The migration of dHL60 during swarming is less organized and less directional compared to primary neutrophils. Isolated neutrophils are loaded on the swarming assay device at a concentration of 2.5 x 10^6^ cells/mL. The time interval between frames is 5 minutes. Scale bar is 50 μm.

**Supplementary Video 5. dHL-60 swarming depends on LTB4-mediated cell-cell communication**. dHL-60 were treated with BLT1 & 2 receptors antagonists, for at least 30 minutes. The nucleus of dHL-60 was stained with Hoechst dye. Treated dHL-60 formed smaller swarms around zymosan particle cluster spots compared to untreated dHL60s. One spot (140 μm in diameter) is identified by a broken red circle. dHL-60 cells are loaded on the device at a concentration of 2.5 x 10^6^ cells/mL. The time interval between frames is 1 minute. Scale bar is 50 μm.

**Supplementary Video 6: dHL-60 swarming depends on LTB4-mediated cell-cell communication**. dHL-60 were treated with MK-886 pathway inhibitor, for at least 30 minutes. The nucleus of dHL-60 was stained with Hoechst dye. Treated dHL-60 formed smaller swarms around zymosan particle cluster spots compared to untreated dHL60s. One spot (140 μm in diameter) is identified by a broken red circle. dHL-60 cells are loaded on the device at a concentration of 2.5 x 10^6^ cells/mL. The time interval between frames is 1 minute. Scale bar is 50 μm.
